# Inductive reasoning differs between taxonomic and thematic contexts: Electrophysiological evidence

**DOI:** 10.1101/333211

**Authors:** Fangfang Liu, Jiahui Han, Lingcong Zhang, Fuhong Li

## Abstract

Inductive reasoning can be performed in different contexts, but it is unclear whether the neural mechanism of inductive reasoning performed in a thematic context (e.g., panda has x, so bamboo has x) is the same as that performed in a taxonomic context (e.g., panda has x, so bear has x). In the present study, participants were required to judge whether a conclusion was acceptable or not based on its premise, for which the taxonomic or thematic distances between premise and conclusion objects were either far or near. The ERP results indicated that the effect of reasoning context (taxonomic vs. thematic) was initially observed in the P2 component; while the distance effect (far vs. near) was observed in N400 and late components. Moreover, the distance effect on thematic-based inductive reasoning was found in the frontal and frontal-central brain regions, while the distance effect in taxonomic-based inductive reasoning conditions was found in the central-parietal and parietal regions. These results support the view that inductive reasoning is performed differently under different semantic contexts.

## Introduction

Inductive reasoning involves collecting and remembering instances of a rule, generating a hypothesis based on these instances, integrating new instances, and confirming the resulting hypothesis through further observation [1]. In recent years, inductive reasoning and its cognitive neural mechanism have widely been studied using various tasks [4, 6, 17, 26, 31-35, 43-44].

A number studies have demonstrated that inductive reasoning is performed in taxonomic-based contexts [1, 33, 35-36, 38-39]. For an instance, Long et al. revealed that, in taxonomic contexts, illogical conclusions evoked greater-amplitude of N2 and N400 than those of logical conclusions.

Other studies have investigated how the thematic relations are processed in inductive reasoning, and compared this to the processing of taxonomic relations [36, 39]. It has been found that the processing of thematic relation requires fewer cognitive resources than the processing of taxonomic relation in reasoning tasks [25, 29, 47, 49]. Taxonomic relation refers to an overlap in the features or meaning of a word. However, thematic relation includes externally or complementarily related items within scenarios or events (e.g., closet and dress, dog and leash, etc.), which shares an associative relationship but not perceptual features [11, 25, 49]. Taxonomic relations seem to be processed semantically by retrieving category knowledge, while thematic relations are more likely to be processed automatically [10, 25]. Alternatively, Kalénine et al. suggested that taxonomic relations rely on perceptual processes while thematic relations rely on event/action processing [24].

As reviewed above, existing studies have primarily adopted imaging methods to explore the difference between taxonomic- and thematic-based inductive reasoning, while the temporal dynamics of brain activation associated with the difference remains unaddressed. The purpose of the present study was to investigate the electrophysiological distinction between these two types of inductive reasoning. Particularly, we tested whether distance effects on the processing of taxonomic- and thematic-based semantic relations in inductive reasoning were differently reflected in brain responses. We designed four conditions, including taxonomic-far, taxonomic-near, thematic-far, and thematic-near conditions, to test this. In each trial, we sequentially presented a premise and a conclusion. Participants had to decide whether the conclusions were acceptable based on the premise or not.

First, we predicted that the difference would initially be observed in an earlier component, such as the P2, which is related to perceptual analysis, visual encoding, and visual attention [4, 35, 44]. Since the processing of taxonomic relations requires more cognitive effort than thematic relationships [29], we hypothesized that taxonomic processing would evoke a larger P2 than would thematic processing. Second, we hypothesized that the distance effect on both types of processing would be observed in the N400 component [35, 38]. We also expected that the scalp distribution of the N400 distance effect would be different between these two types of inductive reasoning, because cerebral cortex activation associated with thematic knowledge has been reported to be different from that of taxonomic knowledge [24, 47, 49].

## Methods

### Participants

Sixty-four healthy undergraduate students (aged 18-25 years) rated the experimental materials in a pilot test. Another 24 healthy undergraduate students (19 female, mean age = 20.03 years, range: 18-24 years, *SD* = 1.45) participated in the formal experiment. All participants were right-handed with normal or corrected-to-normal vision. The experiment was approved by the ethics review board of Jiangxi Normal University.

### Stimuli and experimental procedure

In the pilot experiment, participants were required to rate the strength of thematic and taxonomic relations for each word pair on a 7-point scale (7: very relevant, 4: moderate relevance, 0: completely irrelevant). All word pairs were selected from previous studies and translated into Chinese [4, 15, 25, 40, 53].

Four types of word-pairs were designed, with a total of 256 pairs of words. Sixty-four pairs had a thematic-far relation, whereby the two words were thematically related with a far distance (e.g. panda vs. tree); 64 pairs had a thematic-near relation, whereby the two words were thematically related with a near distance (e.g. panda vs. bamboo); 64 pairs had a taxonomic-far relation, whereby the two words were taxonomically related with a far distance (e.g. panda vs. dolphin); and 64 pairs had taxonomic-near relation, whereby the two words were taxonomically related with a near distance (e.g. panda vs. antelope).

Finally, the 64 pairs of words in the four conditions were retained and used in the formal experiment. A paired-*t* test indicated that the thematic-near word pairs (*M* = 6.08, *SD* = 0.28) were significantly different from the thematic-far word pairs (*M* = 4.39, *SD* = 1.05) in terms of thematic relation, *t* (1, 126) = 12.48, *p* < 0.01. The taxonomic-near word pairs (*M* = 5.62, *SD* = 0.46) differed significantly from the taxonomic-far word pairs (*M* = 3.77, *SD* = 0.86) in terms of taxonomic relation, *t* (1, 126) = 16.78, *p* < 0.01.

The formal experiment was a category-based induction task [38]. Each trial consisted of a premise and a conclusion. At the end of each trial, a question mark indicated that participants should decide whether the conclusion was acceptable or not based on the premise. The premise was a sentence that stated that an object had a novel property (e.g. X1). The novel property was presented by mixing a capital letter with an Arabic number ranging from 1-9 (e.g. X1), which served as the blank property. The blank property was also regarded as a meaningless property, which would reduce the influence of background knowledge on reasoning.

Four types of arguments (Taxonomic-Near, Taxonomic-Far, Thematic-Near, Thematic-Far) were presented sequentially on a computer screen and each argument was repeated twice. The entire formal experiment comprised 512 trials (128 in each condition). As shown in Fig 1, a fixation cross was presented in the center of a black screen for 500 ms at the beginning of each trial. After a blank screen (800–1200 ms), the premise was presented for 500 ms, followed by another blank screen for 800–1200 ms. Next, the conclusion appeared on the screen and remained until participants responded. Participants were instructed to respond to the conclusions as rapidly and accurately as possible. They were asked to press the “F” key for “yes” and the “J” key for “no” using the left or right index finger, respectively. The keys for different responses were counterbalanced across participants. Twenty-five practice trials were completed before the test to familiarize participants to the procedure. The arguments used in practice trials were not included in the formal experiment.

**Fig. 1.**
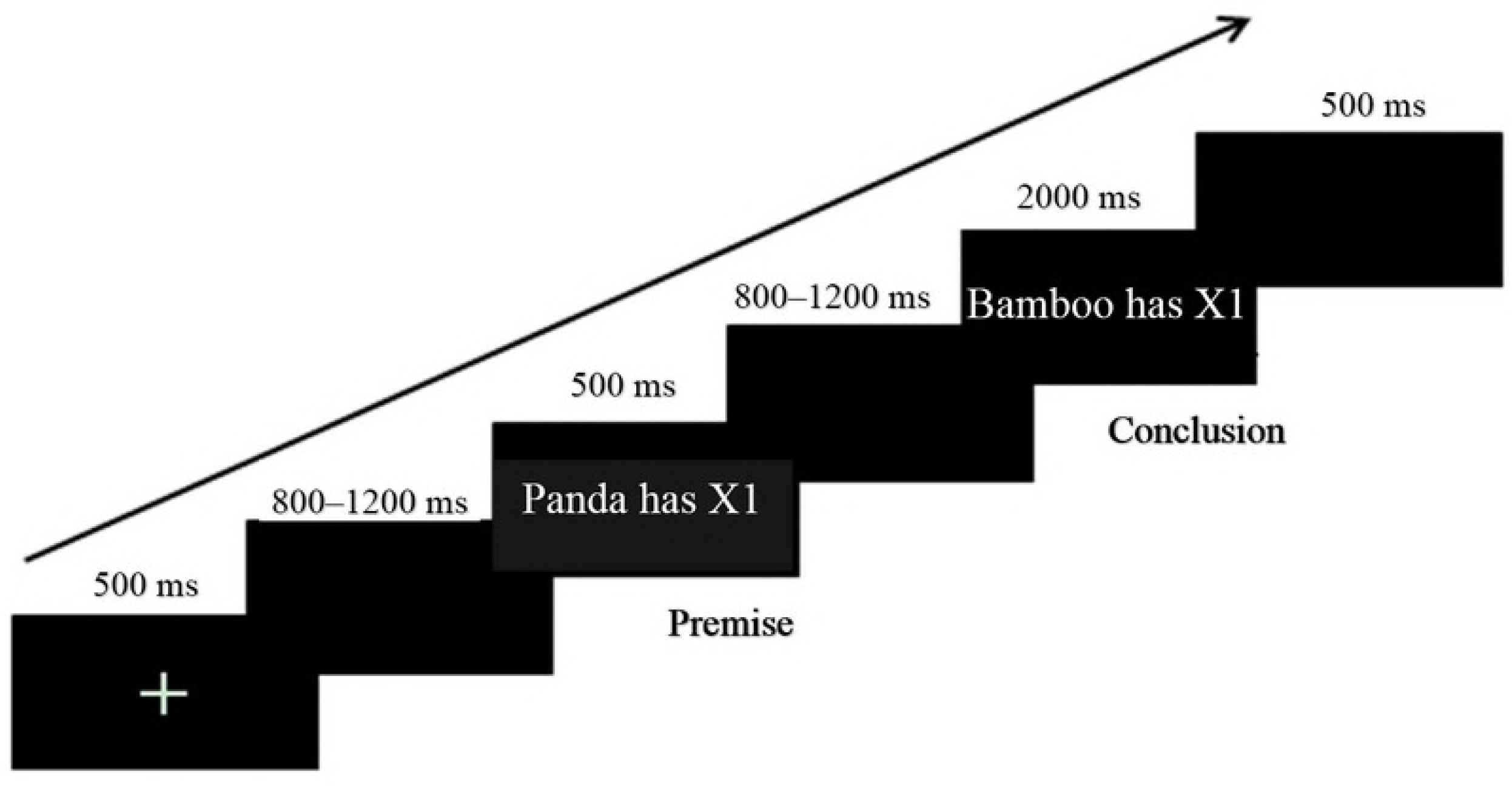
Experimental design and the procedure for a trial.

### ERP recordings and statistical analyses

Electrophysiological activity was recorded using a 64-channel EEG system (Brain Products GmbH, Munich, Germany), with the reference electrodes on the left and right mastoids. An electrode placed under the right eye (for electrooculography; EOG) allowed the monitoring of blinks and vertical eye movements. The impedance of all electrodes was maintained below 5 kΩ. Raw data were band-pass filtered between 0.01–100 Hz and digitized at a sampling rate of 500 Hz. Trials with EOG artifacts (a mean EOG voltage exceeding ±80 μV), and those contaminated with artifacts due to amplifier clipping, bursts of electromyographic activity, or peak-to-peak deflections exceeding ±80 μV were excluded from averaging.

Data were collected continuously and analyzed off-line using Brain Vision Analyzer Software 2.1 (BrainProducts, Munich, Germany). Frequencies lower than 0.1 Hz or higher than 30 Hz were digitally filtered (24 dB). The analysis epoch was 1000 ms with respect to the averaged voltage over the 200-ms epoch before the onset of the conclusion stimulus. The ERP waveforms were time-locked to the onset of the conclusion stimuli. The averaged epoch for the ERPs to the conclusion stimuli, including a 200-ms pre-stimulus baseline, was 1200 ms. According to visual inspection of the grand average waveforms and to previous studies [35, 38], the P2 (190–240 ms), N400 (360–440 ms), and late negative component (LNC) (500–800 ms) at 15 electrode sites (F3, Fz, F4, FC3, FCz, FC4, C3, Cz, C4, CP3, CPz, CP4, P3, Pz, and P4) were analyzed. The mean amplitudes of each component were analyzed using a 2 (distance: far vs. near) × 2 (context: thematic vs. taxonomic) × 3 (laterality: left, middle, right) × 5 (frontality: frontal, frontal-central, central, parietal-central, parietal) repeated measures ANOVA. For all analyses, the *p* values were corrected for deflections according to the Greenhouse-Geisser method.

## Results

### Behavioral Results

Reaction times (RTs) that were extremely large or small (±3 *SD* beyond the mean) were removed. The 2 (context: thematic vs. taxonomic) × 2 (distance: far vs. near) repeated measures analysis of variance (ANOVA) showed a main effect of distance, *F* (1, 28) = 16.54, *p* < 0.001, *η*^*2*^ = 0.37, with longer RTs for far distance. No significant difference was found between the different types (*p* = 0.54). There was no interaction between distance and type (*p* = 0.69).

### ERP Results

### P2 (190–240 ms)

The ERPs evoked by the conclusion in the different conditions are shown in Fig 2. There was an interaction between context and laterality, *F* (2, 46) = 7.55, *p* < 0.01, *η*^*2*^= 0.25. Simple-effect tests showed that taxonomic relations elicited a larger P2 than did thematic relations in the left (*p* = 0.002), middle (*p* < 0.01), and right sites (*p* < 0.01). There was a main effect of context, *F* (1, 23) = 25.96, *p* < 0.01, *η*^*2*^ = 0.53. There was no effect of distance and no other interaction.

**Fig. 2.**
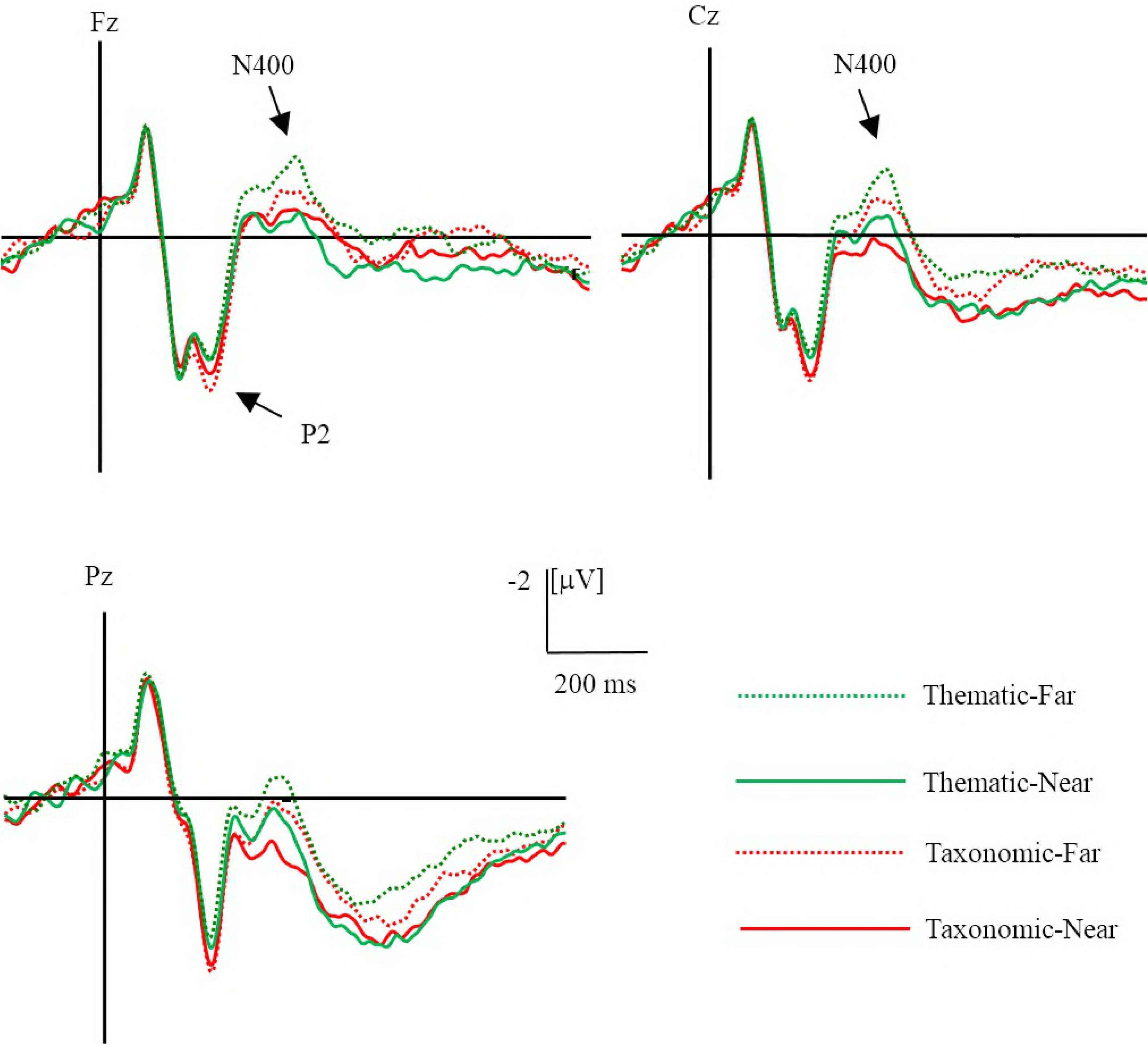
Grand averaged (*n* = 24) ERPs evoked by different conditions.

### N400 (360–440 ms)

A three-way interaction of context, frontality, and laterality was observed, *F* (8, 184) = 3.65, *p* = 0.001, *η*^*2*^ = 0.14. A simple-effect analysis revealed an effect of context at the following sites: FC3, C3, Cz, CPz, Pz, C4, and CP4 (all, *p* < 0.05), with a larger N400 for thematic relations than for taxonomic relations. An interaction between context and frontality was observed, *F* (4, 92) = 4.27, *p* = 0.003, *η*^2^ = 0.16. A simple-effect analysis showed that thematic relations elicited a larger N400 than did taxonomic relations at central (*p* = 0.014) and central-parietal (*p* = 0.015) areas.

There was a three-way interaction of context, distance, and frontality, *F* (4, 92) = 8.37, *p* < 0.01, *η*^*2*^ = 0.27. A simple-effect analysis showed that a distance effect on thematic relation was observed in the frontal (*p* = 0.013) and frontal-central (*p* = 0.011) regions, while the distance effect on taxonomic relation were found in the central-parietal (*p* = 0.001) and parietal (*p* = 0.001) regions (Fig 3). Furthermore, there was a two-way interaction of distance and laterality, *F* (2, 46) = 7.61, *p* = 0.001, *η*^2^ = 0.25. A simple-effect analysis showed that far distance elicited a larger N400 than did near distance conditions at the left (*p* = 0.005), middle (*p* = 0.001), and right sites (*p* = 0.002).

**Fig. 3.**
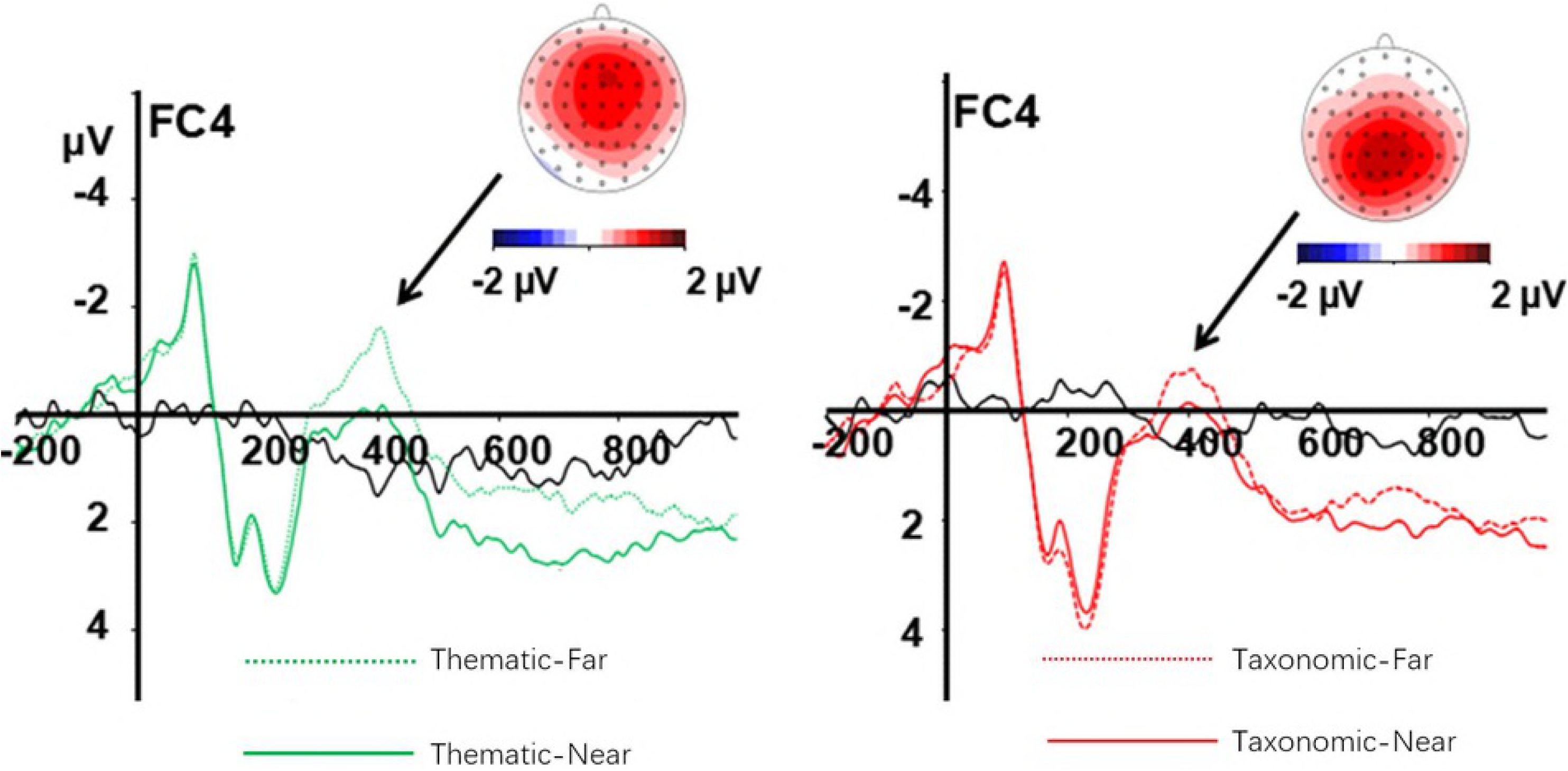
Difference waves and topographical maps of the distance effect on the N400 for thematic (left) and taxonomic (right) conditions.

There was a main effect of context, *F* (1, 23) = 5.02, *p* = 0.035, *η*^*2*^ = 0.18, and effect of distance, *F* (1, 23) = 12.57, *p* = 0.002, *η*^*2*^ = 0.35. Thematic relations significantly elicited a larger N400 amplitude than taxonomic relations, and the far distance generally elicited a larger N400 amplitude than near distance.

### LNC (500–800 ms)

To investigate the time course of the LNC more precisely, we analyzed three successive intervals, 500**–**600 ms, 600**–**700 ms, and 700**–**800 ms. Statistical analysis revealed a main effect of distance in all LNC latency windows (all *p* < 0.01). There was an interaction between distance and frontality in each latency window, *F*_500-600 ms_ (4, 92) = 3.64, *p* = 0.008, *η*^2^ = 0.14; *F*_600-700 ms_ (4, 92) = 2.83, *p* = 0.029, *η*^2^ = 0.11; *F*_700-800 ms_ (4, 92) = 2.59, *p* = 0.042, *η*^2^ = 0.10. Simple-effects tests showed that far distance evoked more negative waves than did near distance conditions in all regions within each time window, 500**-**600 ms: *p*_frontocentral_ = 0.031, *p*_central_ = 0.004, *p*_parietocentral_ = 0.002, and *p*_parietal_ = 0.003; 600**-**700 ms: *p*_frontocentral_ = 0.013, *p*_central_ = 0.003, *p*_parietocentral_ = 0.001, and *p*_parietal_ < 0.001); 700**-**800 ms: *p*_frontocentral_ = 0.009, *p*_central_ = 0.001, *p*_parietocentral_ = 0.001, and *p*_parietal_ < 0.001.

Within the 600**–**700 ms time window, an interaction of distance and laterality was found, *F* (2, 46) = 4.23, *p* = 0.021, *η*^2^ = 0.16. A simple-effect analysis showed that far distance evoked greater negative waves than near distance conditions in the left (*p* = 0.018), middle (*p* = 0.002), and right sites (*p* = 0.003).

## Discussion

The main purpose of this study was to differentiate the electrophysiological response to inductive reasoning under thematic and taxonomic contexts. The behavioral results showed that RTs are significantly longer in far distance than in near distance conditions, regardless of the context type, which indicates that the reasoning process between premise and conclusion required more effort in far distance than near distance conditions [12, 19, 46, 58].

ERP results revealed the effect of experimental condition in three time windows, corresponding to the P2, N400, and LNC components. In the P2 time window, there was an effect of context (thematic vs. taxonomic), but no effect of distance (near vs. far). During the N400 time window, both the context effect and distance effect were observed. In the LNC time window, only a distance effect was observed.

Consistent with our predictions, the difference between the two contexts of inductive reasoning were initially observed in the P2 time window. Taxonomic-based inductive reasoning elicited a larger P2 amplitude than thematic trials. The P2 is generally associated with allocation of attention [2, 7, 42], perceptual encoding [14, 30, 35, 55], and early semantic processes [28, 40]. In the present study, the effect of context on the P2 component might be related to the earlier process of relation encoding [30]. The formation of taxonomic relations is based on an overlap in features of category members. The closest connection between pandas and lions is that they are both terrestrial mammals. With regards to pandas and dolphins, even though they are both mammals, pandas live on land and walk on four legs, while dolphins live in water and have no legs. Therefore, it is necessary to compare common perceptual traits and other major behavioral characteristics (such as breast-feeding) between two species to find the taxonomic relationship between them [11, 40, 47, 49]. In contrast, when looking for the thematic relation between two species, participants do not compare the perceptual characteristics, but simply remember whether there is a thematic relationship between them. That is, the establishment of the relationship between a panda and bamboo only requires the knowledge that a panda eats bamboo. Therefore, compared to the processing of thematic relations, the early process of taxonomic relations evoked a greater P2 amplitude.

It is necessary to note that there was only a context effect, but no distance effect on the P2 component. This illustrated that, in an early time window such as that of the P2 component, participants distinguish relation types before proceeding to the next stage, semantic processing, which was associated with the N400.

In the N400 time window, both a context effect (thematic vs. taxonomic) and distance effect (near vs. far) were observed. This indicates that, after encoding the related feature of two categories (premise and conclusion) within the P2 time window, participants made an elaborative semantic integration of the relation between premise object and conclusion object, and determined whether the conclusion object had the same property as the premise object.

The N400 component is typically related to semantic integration [38, 54] and semantic anomalies [45]. For example, incoherent words or sentences evoke larger-amplitude N400 components [10, 13, 20, 47-48, 20]. The N400 has also been observed in previous ERP studies on category-based inductive reasoning [38, 55]. During reasoning, the concept is the basic unit, and humans principally conduct reasoning using conceptual information [15]. Two concepts do not form a semantic relation until they have an intersection. Semantic distance is an influencing factor for inferring with the degree of relation between two concepts [21]. Kmiecik and Morrison investigated verbal analogical reasoning with different semantic distances, and found that near analogical distance elicits less negative N400 components than does far analogical reasoning [27].

In the present study, we used an inductive reasoning task and manipulated the context and distance between premise and conclusion. We found that, for both taxonomic and thematic conditions, far relation elicited larger N400 amplitudes than near relation reasoning. This result is consistent with existing studies on analogical reasoning, which suggests that semantic distance has a significant effect on reasoning within the N400 time window [19, 46]. That is, in near distance conditions, it is easier to integrate and infer semantic relations between a premise and a conclusion. In contrast, for far distance, it is difficult to identify the intersection between two concept nodes, which evokes a larger N400 amplitude than that of near distance conditions.

Although distance effects were observed in the N400 for both two types of reasoning, the distance effect on thematic-based reasoning was mainly observed in the frontal and frontal-central brain regions, while distance effect on taxonomic-based reasoning was observed in the central-parietal and parietal region. Previous studies have shown that inductive reasoning mainly involves the left medial frontal or the left frontal gyrus [17, 38] and an effect of semantic distance on analogical reasoning was found in the left frontopolar cortex [17, 19, 38]. Our finding is partially consistent with these studies, and indicates that the processing of thematic relationships is associated with activation of the prefrontal cortex [10]. However, taxonomic-based reasoning involved the process of comparing and analyzing critical or typical features of the objects in the premise and conclusion. Previous studies have shown that the comparison of perceptual characteristics is primarily associated with activation of the parietal cortex [3].

In conclusion, we used a modified inductive reasoning task to investigate the electrophysiological difference between thematic and taxonomic-based inductive reasoning. ERP results revealed a significant effect of distance on both types of reasoning during the N400 time window, but the scalp distribution of this distance effect was different between these two types of semantic processing, with a frontal distribution for thematic-based reasoning and a posterior distribution for taxonomic-based reasoning. This supports the view that these two types of semantic processing might recruit different neural networks.

